# Protease OMA1 Modulates Mitochondrial Bioenergetics and Ultrastructure through Dynamic Association with MICOS Complex

**DOI:** 10.1101/2020.11.24.395111

**Authors:** Martonio Ponte Viana, Roman M. Levytskyy, Ruchika Anand, Andreas S. Reichert, Oleh Khalimonchuk

## Abstract

Remodeling of mitochondrial ultrastructure is a complex dynamic process that is critical for a variety of mitochondrial functions and apoptosis. Although the key regulators of this process - mitochondrial contact site and cristae junction organizing system (MICOS) and GTPase Optic Atrophy 1 (OPA1) have been characterized, the mechanisms behind this regulation remain incompletely defined. Here, we found that in addition to its role in mitochondrial division, metallopeptidase OMA1 is required for maintenance of contacts between the inner and outer membranes through a dynamic association with MICOS. This association is independent of OPA1, appears to be mediated via the MICOS subunit MIC60, and is important for stability of MICOS machinery and the inner-outer mitochondrial membrane contacts. We find that OMA1-MICOS relay is required for stability of respiratory supercomplexes, optimal bioenergetic output in response to cellular insults, and apoptosis. Loss of OMA1 affects these activities; remarkably it can be partially compensated for by an artificial MICOS-emulating tether protein that bridges the inner and outer mitochondrial membranes. Our data show that OMA1-mediated support of mitochondrial ultrastructure is required for maintenance of mitochondrial architecture and bioenergetics under both basal and homeostasis-challenging conditions, and suggest a previously unrecognized role for this protease in mitochondrial physiology.

## INTRODUCTION

Mitochondria are dynamic organelles with distinctive and complex architecture, wherein two disparate membranes, termed the outer mitochondrial membrane (OM) and the inner mitochondrial membrane (IM), form specific sub-compartments – the mitochondrial matrix and the intermembrane space (IMS). Unlike the OM, the IM is topologically heterogeneous and comprises two distinct regions: the inner boundary membrane (IBM), a planar membrane located in the vicinity of the OM, and the inner cristae membrane (ICM) – the region of IM that forms large invaginations termed cristae (Frey and Manella, 2000; Zick et al., 2009). These tubular- or lamellar-like structures are highly enriched in respiratory complexes and stabilized by structures known as cristae junctions (Glickerson et al., 2003; Zick et al., 2009; Cogliati et al., 2016). Such architecture is at least in part mediated by a large protein complex known as MICOS (for mitochondrial contact site and cristae junction organizing system), which has been shown to have a crucial role in the formation and maintenance of cristae junctions and OM-IM contacts (Rabl et al., 2009; Harner et al., 2011; Hoppins et al., 2011; van der Laan et al., 2016; Wollweber et al., 2017; Pfanner et al., 2014). MICOS resides in cristae junctions and encompasses at least six evolutionary conserved subunits: MIC60/Mitofilin, MIC19/CHCHD3, MIC25/CHCHD6, MIC27/APOOL, MIC26/APOO, and MIC10/MINOS1 (van der Laan et al., 2016). Other factors important for cristae ultrastructure include F1FO-ATP synthase (Davies et al., 2012; Daum et al., 2013) and Optic Atrophy 1, OPA1 (Frezza et al., 2006; Yamaguchi et al., 2008; Cogliati et al., 2013; Patten et al., 2014; Varanita et al., 2015). The latter is an IM-resident dynamin-like GTPase related to OM-associated GTP hydrolases DRP1 and Mitofusins 1 and 2, which have been shown to reciprocally regulate division and fusion of the mitochondrial network, respectively (Smirnova et al., 2001; Yoon et al., 2001; Chen et al., 2003; Song et al., 2009). OPA1 exists as a nearly stoichiometric mix of IM-anchored long (L-OPA1) and soluble IMS-localized short (S-OPA1) forms (Duvezin-Caubet et al., 2006; Ishihara et al., 2006). In addition to its well-established role as a pro-fusion factor in the maintenance of the mitochondrial network (Ishihara et al., 2006; Song et al., 2009; MacVicar and Langer, 2016), OPA1 variants form oligomers that are believed to connect cristae junctions, thereby segregating cristae content from the IMS and stabilizing respiratory supercomplexes (Cogliati et al., 2013; Patten et al., 2014; Yoon et al., 2017). Interestingly, OPA1 and MICOS oligomers appear to be in physical contact and have been proposed to regulate cristae shape in a semi-cooperative manner (Darshi et al., 2011; Barrera et al., 2016; Schweppe et al. 2017; Glytsou et al., 2016;).

Changes in cellular metabolic demands or homeostatic insults are known to correlate with changes in the mitochondrial network and ultrastructure (Hackenbrock 1966; Cogliati et al., 2013; Patten et al., 2014; Plecita-Hlavata et al., 2016). A number of studies implicated OPA1 as a central component behind these alterations (Cogliati et al., 2013; Mishra et al., 2014; Patten et al., 2014; Zhang et al., 2011). Although the significance of the L- and S-OPA1 variants in these events remains debated (Mishra et al., 2014; Anand et al., 2014; Lee et al., 2017), the OPA1-dependent IM remodeling is recognized as pivotal to several vital mitochondrial functions such as energy conversion and apoptosis (Frezza et al., 2006; Yamaguchi et al., 2008; Tondera et al., 2009; Mishra et al., 2014; Anand et al., 2014; Korwitz et al., 2016). Two IM proteases, the i-AAA protease YME1L and the stress-activated peptidase OMA1 are critical regulators of IM dynamics in higher eukaryotes (Rainbolt et al., 2016; MacVicar and Langer, 2016; Levytskyy et al., 2017). They modulate the mitochondrial network via proteolytic processing of L-OPA1 variants under basal and stress conditions, respectively (Head et al., 2009; Ehses et al., 2009; Baker et al., 2014; Zhang et al., 2014; Anand et al., 2014; Wai et al., 2016). In the latter case, rapid proteolytic conversion of all available L-OPA1 to the S-OPA1 form by OMA1 promotes fragmentation of the mitochondrial network – a critical event required for several downstream mechanisms such as apoptosis or mitophagy (Frezza et al., 2006; Yamaguchi et al., 2008; Rambold et al., 2011; Anand et al., 2014; MacVicar and Lane, 2014). As such, preservation of L-OPA1 variant through its overexpression or OMA1 depletion has been shown to stabilize the mitochondrial network, maintain normal cristae structure and exert anti-apoptotic effects (Anand et al., 2014; Varanita et al., 2015). On the other hand – despite normal mitochondrial ultrastructure – OMA1-deficient mouse embryonic fibroblasts (MEFs) are bioenergetically compromised under conditions that demand maximal respiratory output (Quiros et al., 2012; Korwitz et al., 2016; Bohovych et al., 2015). OMA1 is also known to perform protein quality control functions and mediate the stability of respiratory supercomplexes, thus influencing mitochondrial energy metabolism (Desmurs et al., 2015; Bohovych et al., 2015). An important unresolved question regards mechanistic aspects and the functional connection of these activities.

In the present study, we show that in addition to its role in mitochondrial fission, OMA1 is required for the organelle&#8217;s ultrastructural organization through its association with MICOS machinery. This interaction is likely mediated by the MIC60-MIC19 subcomplex of MICOS and influences the recruitment of other MICOS proteins without significantly impacting mitochondrial fission/fusion machinery. We show that OMA1-mediated MICOS stabilization is particularly important for optimal mitochondrial performance in response to cellular insults such as mitochondrial depolarization. Remarkably, in the absence of OMA1, these activities can be partially compensated for by an artificial tether protein that bridges the OM and IM, thereby emulating the tunable MICOS complex.

Our results demonstrate that OMA1-mediated support of mitochondrial intermembrane contacts is required for modulation of mitochondrial bioenergetic in response to homeostatic challenges and suggest a previously unrecognized role for this protease in mitochondrial physiology.

## RESULTS

### OMA1 associates with MIC60 component of MICOS complex

In our effort to gain new insight into the physical interactome of OMA1, we screened for its physiologically relevant interacting partners using the recently developed BioID method (Roux et al., 2013). This non-biased approach involves fusing the protein of interest to promiscuous BirA biotin ligase, which specifically modifies proteins in the immediate vicinity to the bait. HEK293T cells expressing OMA1-BirA or a topologically matched SURF1-BirA fusion construct (non-specific binding control) were cultured to allow the modification, lysed and subjected to affinity purification of biotinylated proteins followed by their identification by LC-MS/MS (Supplementary Fig. S1A). Subsequent analysis of candidate hits eliminated known contaminants and non-specific interactors. One prey protein that withstood our elimination analysis was Mitofilin/MIC60, a core component of the MICOS complex. To validate this interaction, we used HEK293T cells expressing C-terminally Myc-tagged MIC60. Immunoprecipitation of epitope-tagged MIC60 also led to specific co-purification of endogenous MIC60 and OMA1, but not the abundant m-AAA protease subunit AFG3L2 (Fig. 1A). We have also detected some amounts of OPA1, which has been previously reported to interact with MIC60 (Glytsou et al., 2016; Barrera et al., 2016). Reciprocally, endogenous MIC60 co-precipitated with the GFP-tagged OMA1 (Supplementary Fig. S1B). Consistent with these results, OMA1 and MIC60 high molecular weight pools co-fractionated on sucrose density gradient (Fig. 1B). We also observed partial co-fractionation of these proteins with OPA1 signal.

**Figure 1.**
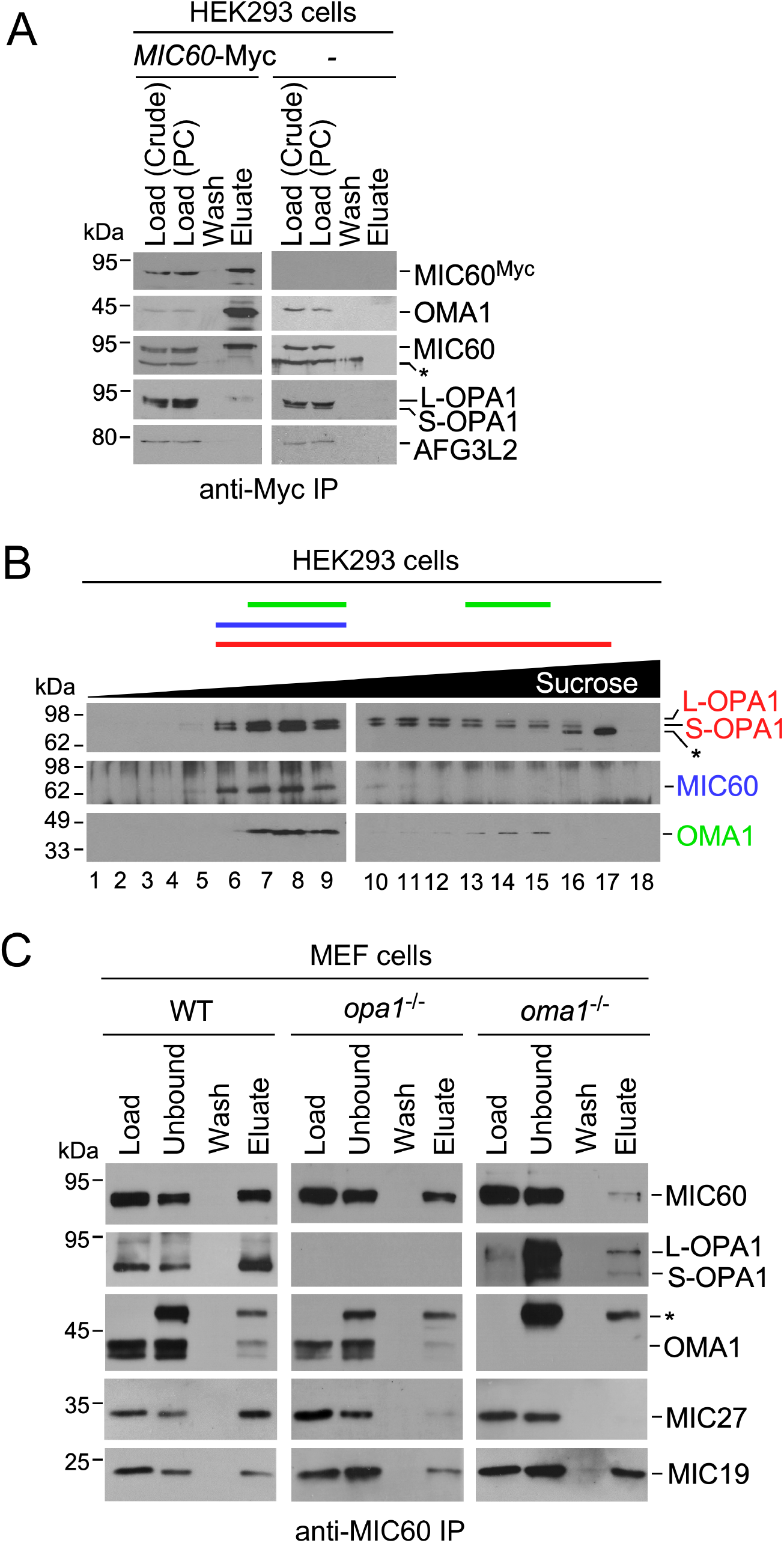
Evolutionarily conserved interaction between OMA1 and MIC60. **(A)** HEK293T cells were transfected with MIC60-Myc construct or mock transfected. Mitochondrial lysates from the respective cells were pre-cleared with IgG beads and then immunoprecipitated with anti-Myc magnetic beads. Precipitated proteins from the initial sample [load (crude)], the sample following pre-clearing [load (PC)], the last wash, and the eluate were analyzed by western blot with indicated antibodies. The asterisk indicates a non-specific band. **(B)** Digitonin-solubilized mitochondria from HEK293T cells were fractionated in a 12-50% sucrose density gradient by centrifugation. Collected fractions were analyzed by immunoblotting with antibodies against indicated proteins. The asterisk indicates a non-specific band. **(C)** Mitochondrial lysates from indicated MEF cell lines were incubated with agarose beads-coupled anti-MIC60 antibodies. Immunoprecipitated proteins from the initial sample (load), unbound fraction (unbound), last wash (wash), and eluate were analyzed by western blot with indicated antibodies. The asterisk indicates a non-specific band.

Because OPA1 is the substrate of OMA1, and OPA1 and MIC60 are known to interact physically, we wondered if OPA1 might be mediating OMA1-MIC60 interaction. To better understand how these proteins are interrelated, we carried out MICOS co-purifications in mitochondrial lysates from wild type (WT), *opa1*^−/−^ and *oma1*^−/−^ MEFs. Similar to HEK293T mitochondria, immunosorbtion of MIC60 resulted in co-precipitation of both OMA1 and OPA1 along with MIC19, MIC10 and MIC27 MICOS subunits in WT mitochondrial lysates (Fig. 1C and data not shown). Importantly, OMA1-MIC60 interaction remained largely unperturbed in *opa1*^−/−^ mitochondria, indicating that OPA1 is dispensable for OMA1 association with MICOS. Similarly, we found that OPA1 retained its ability to interact with MIC60 in *oma1*^−/−^ cells (Fig.1C). We also noted that loss of OMA1, and to a lesser extent OPA1, negatively impacted MICOS as reflected by impaired MIC60-MIC27 interaction in both *opa1*^−/−^ and *oma1*^−/−^ mitochondria. We conclude that OMA1 is associated with MICOS machinery independently of OPA1.

### Loss of OMA1 impacts mitochondrial structure upon uncoupling

Our finding that OMA1 is physically associated with MICOS prompted us to revisit mitochondrial morphology and ultrastructure in OMA1-deficient cells. We first examined mitochondrial network morphology in WT and *oma1*^−/−^ MEFs. Consistent with previous reports (Quiros et al., 2012; Anand et al., 2014; Korwitz et al., 2016), deletion of OMA1 did not significantly affect mitochondrial tubulation under basal conditions (Fig. 2A). We then assessed the morphology of the mitochondrial network under mild uncoupling – the condition upon which *oma1*^−/−^ MEFs are known to exhibit bioenergetic deficit (Bohovych et al., 2015). As expected under this condition, control MEFs showed rapid fragmentation of mitochondrial network, reflecting OMA1-mediated proteolysis of L-OPA1. The *oma1*^−/−^ cells predictably retained tubular mitochondria upon uncoupling. We noticed, however, that the mitochondrial network morphology was different from that observed in unstressed *oma1*^−/−^ MEFs. Mitochondrial tubules were shorter and tended to form clump-like clusters in CCCP-treated *oma1*^−/−^ cells (Fig. 2A). Steady-state levels of the OM fusion-and fission-mediating proteins were unchanged in these cells (Fig. 2B). As expected, in contrast to the WT cells, the CCCP-treated *oma1*^−/−^ cells retained the long form of OPA1 (Fig. 2C). We next assessed the behavior of the IM GTPase OPA1 by two-dimensional BN-PAGE and observed that uncoupled *oma1*^−/−^ MEFs, unlike WT cells, partially retained high-mass OPA1 species suggesting these oligomers are primarily formed by the L-OPA1 (Fig. 2D).

**Figure 2.**
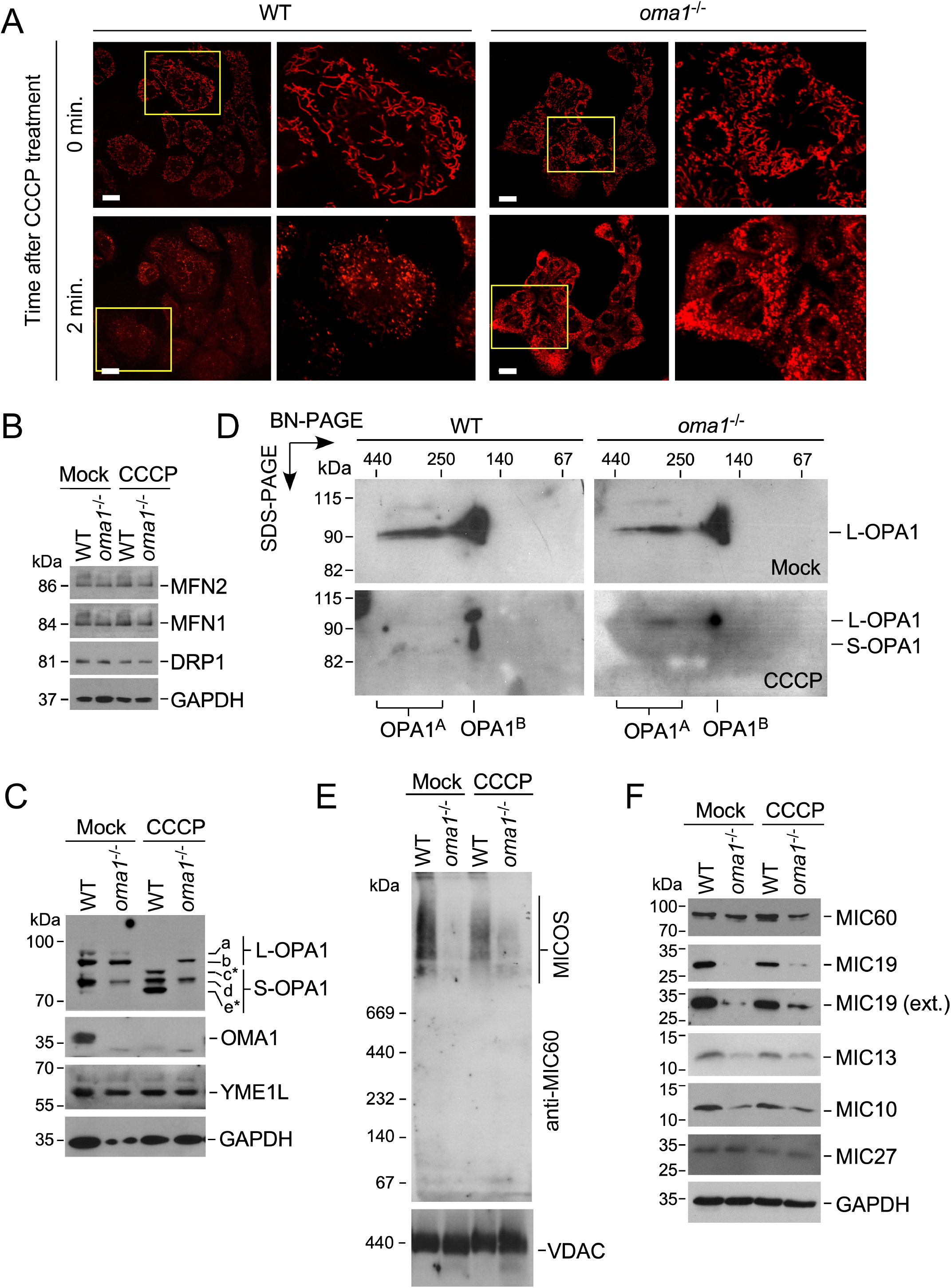
Mitochondrial morphology in normal and stressed *oma1*^*−/−*^ cells. **(A)** *In vivo* confocal microscopy of the mitochondrial network in wild type (WT) and *oma1*^−/−^ MEF cells. Fibroblasts were treated with 2 μM CCCP and incubated for indicated times to induce mild uncoupling, stained with the mitochondrial dye Mitotracker Deep Red and analyzed by confocal fluorescent microscopy. Representative images are shown. Scale bars, 20 μm. **(B, C)** Immunoblot analysis with indicated antibodies of cell lysates from the same samples as in A. In panel C, letters *a-e* denote OPA1 isoforms, with *a* and *b* representing the long form of OPA1, L-OPA1 and *c-e* representing the short form, S-OPA1. Asterisks denote S-OPA1 forms generated via OMA1-mediated cleavage of L-OPA1. **(D)** Two-dimensional native gel electrophoresis analysis of OPA1 oligomers in mitochondria from WT and *oma1*^−/−^ cells treated with (CCCP) or without (Mock) CCCP. OPA1A and OPA1B denote high- and lower-mass OPA1 oligomeric species, respectively. **(E)** Mitochondrial lysates were subjected to one-dimensional blue native gel electrophoresis and analyzed by western blot with antibodies against MIC60 and the outer mitochondrial membrane protein VDAC (loading control). **(F)** Steady-state levels of indicated proteins in mitochondrial lysates from cells described above, analyzed by western blot.

We next sought to test if these changes may be related to OMA1-MIC60 interaction. We examined stability of MICOS high-mass complexes and steady-state levels of key MICOS subunits in cells lacking OMA1. Surprisingly, BN-PAGE analysis revealed marked attenuation of high-mass MICOS complex in *oma1*^−/−^ cells, irrespective of CCCP treatment (Fig. 2E and Supplementary Fig. S2), indicating that MICOS may be more labile under blue-native gel conditions in the absence of OMA1. Consistent with this notion, *oma1*^−/−^ cells exhibited increased accumulation of a lower-mass MIC27-containing complex (Supplementary Fig. S2B), suggesting that the MIC60-MIC19 subcomplex is likely affected by OMA1 loss. We then examined steady-state levels of key MICOS subunits and observed a slight decrease in the levels of MIC60 in *oma1*^−/−^ MEFs under both basal and stress conditions, which suggests this protein is likely not a substrate of OMA1 (Fig. 2F). The steady-state levels of MIC27 subunit were unaltered. By contrast, steady-state levels of MICOS subunits MIC19, MIC13 and MIC10 were decreased in *oma1*^−/−^ cells relative to WT control in both basal and CCCP-uncoupled conditions (Fig. 2F).

We hypothesized that such an impediment may be affecting the architecture of IBM and/or IBM-OM contact sites (Supplementary Fig. S2A), thereby contributing to the altered mitochondrial morphology seen in stressed *oma1*^−/−^ cells (Fig. 3A). To test this postulate, we examined mitochondrial ultrastructure in normal and CCCP-stressed *oma1*^−/−^ MEFs by transmission electron microscopy (TEM). In line with previous reports (Quiros et al., 2012; Anand et al., 2014; Korwitz et al., 2016), we found that about half of unstressed *oma1*^−/−^ cells displayed mitochondrial ultrastructure similar to that of WT MEFs. Interestingly, however, in our hands a significant fraction of analyzed *oma1*^−/−^ MEFs (~51%) exhibited defective cristae ultrastructure (Fig. 3B). Of note, we also observed a small fraction (~14%) of WT cells exhibiting altered cristae morphology – likely due to sample preparation and sectioning conditions. Furthermore, TEM analysis of CCCP-treated *oma1*^−/−^ MEFs revealed that unlike in WT, ~70% of mitochondria in OMA1-deleted cells displayed defective cristae ultrastructure upon uncoupling (Fig. 3C). One particularly pronounced alteration included the appearance of onion-like cristae that were fully detached from the OM. Such ultrastructural changes are comparable to those described in cells depleted for MICOS subunits (Supplementary Fig. S3, Anand et al., 2016; Glytsou et al., 2016) or harboring catalytically dead OPA1 (Lee et al., 2017) and are consistent with our predicted model outlined in Fig. 3A. Altogether, these results suggest that OMA1-MICOS association is required for cristae positioning and/or remodeling, particularly during homeostatic insults.

**Figure 3.**
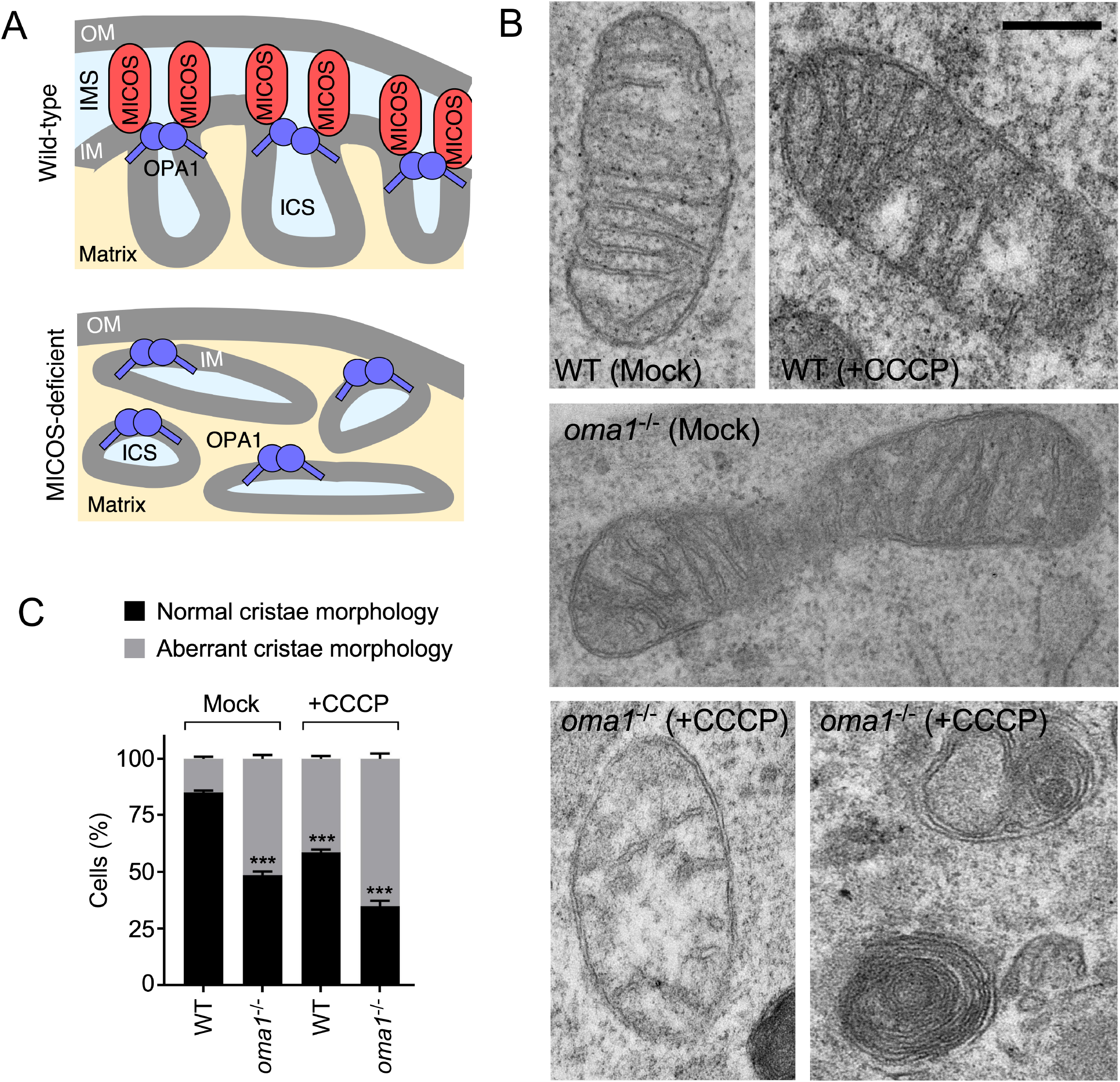
Stressed *oma1*^*−/−*^ cells exhibit altered mitochondrial ultrastructure. **(A)** Schematic depiction of the predicted ultrastructure defect in MICOS-deficient cells. A situation wherein mitochondrial cristae junctions remain closed but the IM-OM contacts are impeded would result in the appearance of vesicle-like cristae detached from the OM. **(B)** Transmission electron microscopy imaging of mitochondrial ultrastructure in wild type (WT) and *oma1*^−/−^ MEF cells incubated with (+CCCP) or without (Mock) 2 μM CCCP. Representative images are shown. Scale bars, 0.4 μm. **(C)** The number of cells bearing mitochondria with aberrant cristae from B was quantified. Error bars, mean ± S.D. of n=3 biological replicates with 100 cells per replicate; \*\*\**p*<0.001 by *t*-test compared to WT without CCCP.

### Emulation of OM-IM contact sites preserves mitochondrial architecture in stressed *oma1*^−/−^ cells

Our data suggest that integrity of OM-IM contacts may be critical for stress-induced cristae remodeling. To validate this idea, we engineered a chimeric construct comprising the N-terminal portion of the IM protein SCO1 (residues 1-161), an unstructured 12 amino acids-long linker region and the transmembrane segment of the OM protein TOMM20 (residues 1-24), followed by a GFP tag (Fig. 4A). This construct, which we called mitoT, is designed to tether the outer and inner mitochondrial membranes, thereby emulating the structural function of MICOS at the IBM sites. Indeed, a similar chimeric tether has been previously shown to partially compensate for the loss of MICOS in yeast (Aaltonen et al., 2016). We generated stable *oma1*^−/−^ MEFs expressing mitoT at levels that were well tolerated by the cells in question (Fig. 4B and C) and showed that this construct was exclusively localized to mitochondria (Fig. 4B). The expression of mitoT had no appreciable effect on survival or proliferation of *oma1*^−/−^ cells, nor did it influence L-OPA1 processing (Supplementary Fig. S4A). Of note, in WT, but not in *oma1*^−/−^ cells, the mitoT protein displayed a slightly faster migration pattern, suggesting that OMA1 might be involved in its processing (Supplementary Fig. S4B).

Next, we examined the ultrastructure of *oma1*^−/−^ [mitoT] cells under basal and stress conditions by TEM. MitoT expression resulted in slightly denser cristae in unstressed cells, relative to *oma1*^−/−^ MEFs (Fig. 4D). More importantly, we observed fewer disorganized cristae: 18% in *oma1*^−/−^ [mitoT] versus 50% in *oma1*^−/−^ MEFs under basal conditions and 45% in *oma1*^−/−^ [mitoT] versus 70% in *oma1*^−/−^ MEFs in CCCP-treated cells (Fig. 4D and E). These data indicate that mitoT is at least in part able to maintain OM-IM contacts in OMA1-deficient mitochondria. Biochemically, the expression of mitoT in *oma1*^−/−^ cells stabilized the high-mass MICOS complex (Fig. 4F) and restored normal steady-state levels of MIC60, MIC19 and MIC10 MICOS subunits (Fig. 4G), further suggesting that mitoT-mediated permanent contacts between the IM and OM result in stabilization of mitochondrial architecture in OMA1-deficient cells.

**Figure 4.**
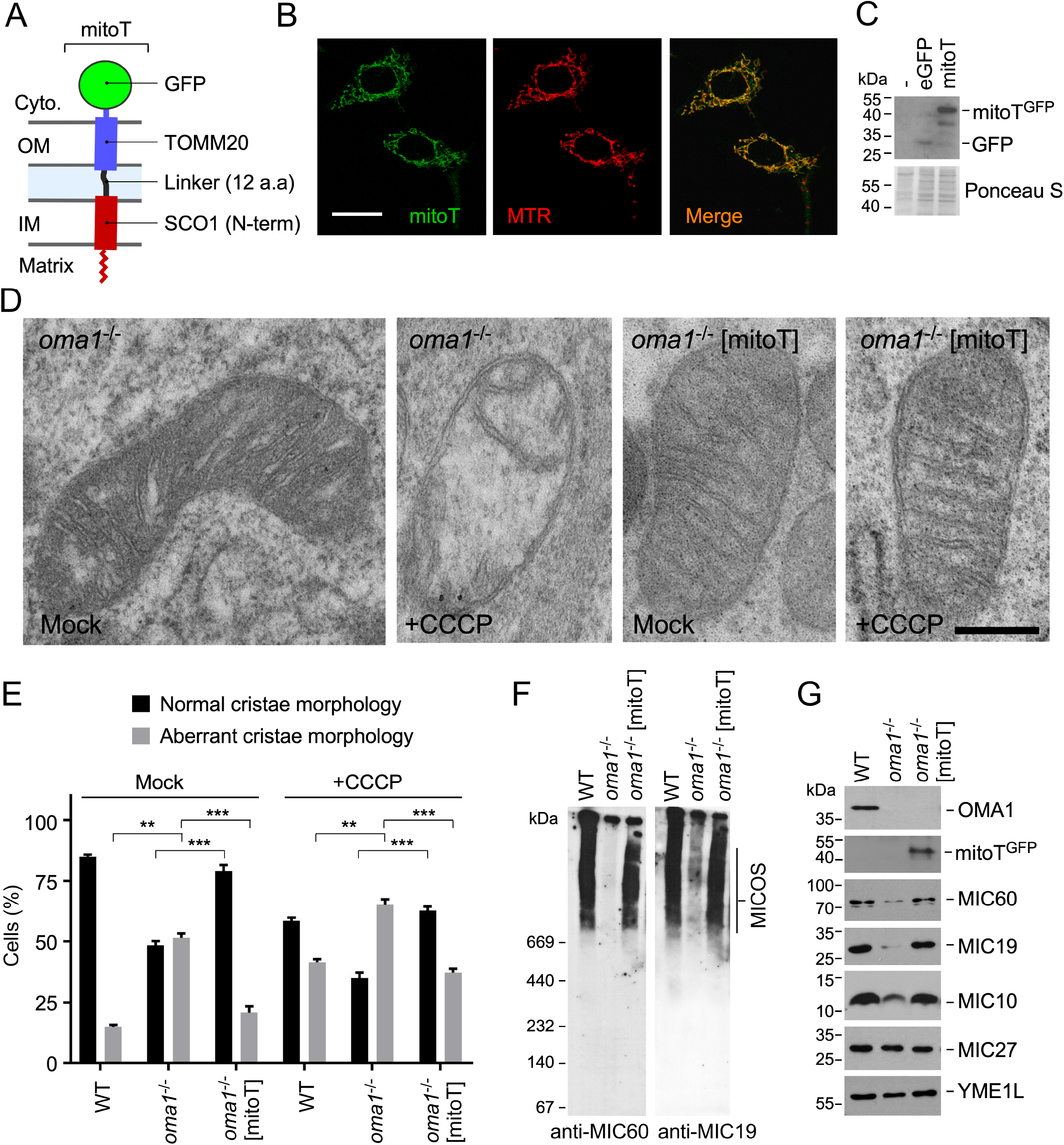
Mitochondrial membranes-bridging chimera partially corrects ultrastructure defects in stressed *oma1*^*−/−*^ cells. **(A)** Schematic depiction of the mitoT artificial tether construct. **(B)***In vivo* confocal microscopy of the mitochondrial network and mitoT-GFP signal in *oma1*^−/−^ MEF cells stably expressing mitoT construct. Representative images are shown for the GFP signal from the tagged mitoT construct (mitoT) and Mitotracker DeepRed (MTR). Scale bar, 20 μm. **(C)** Mitochondrial lysates from cells expressing GFP-tagged mioT and control cells were analyzed by western blotting with anti-GFP antibodies. **(D)** Representative TEM micrographs of mitochondria from control and CCCP-treated *oma1*^−/−^ MEF cells expressing mitoT. Scale bar, 0.4 μm. **(E)** The number of cells bearing mitochondria with normal and aberrant cristae from indicated cell lines was quantified. Error bars, mean ± S.D. of n=3 biological replicates with 100 cells per replicate; \*\**p*<0.01, \*\*\**p*<0.001 by *t*-test. **(F)** Mitochondrial lysates from indicated cell lines were analyzed by BN-PAGE and immunoblotting with antibodies against MICOS subunits MIC60 and MIC19. **(G)** Steady-state levels of indicated proteins in the respective cell lines analyzed by western blot.

### MitoT-mediated OM-IM contacts stabilize respiratory complexes and alleviate bioenergetic deficit in *oma1*^−/−^ cells

Mitochondrial ultrastructure is important for the stability and optimal performance of respiratory supercomplexes (RSCs) (Cogliati et al., 2013; Lee et al., 2017). We therefore wondered if the impaired stability of RSCs and bioenergetic deficit seen in *oma1*^−/−^ cells (Bohovych et al., 2015) could be a consequence of impaired OM-IM connectivity. To test this scenario, we examined mitochondrial lysates from WT, *oma1*^−/−^ and *oma1*^−/−^ [mitoT] MEFs by blue-native gel electrophoresis. Consistent with our previous observations, the abundance and stability of higher order RSCs was slightly decreased in *oma1*^−/−^ MEFs under the conditions tested (Fig. 5A). By contrast, we found that mitoT expression stabilized the RSCs in these cells (Fig. 5A). Steady-state levels of the representative complex subunits remained unchanged in all cell lines tested (Fig. 5B).

**Figure 5.**
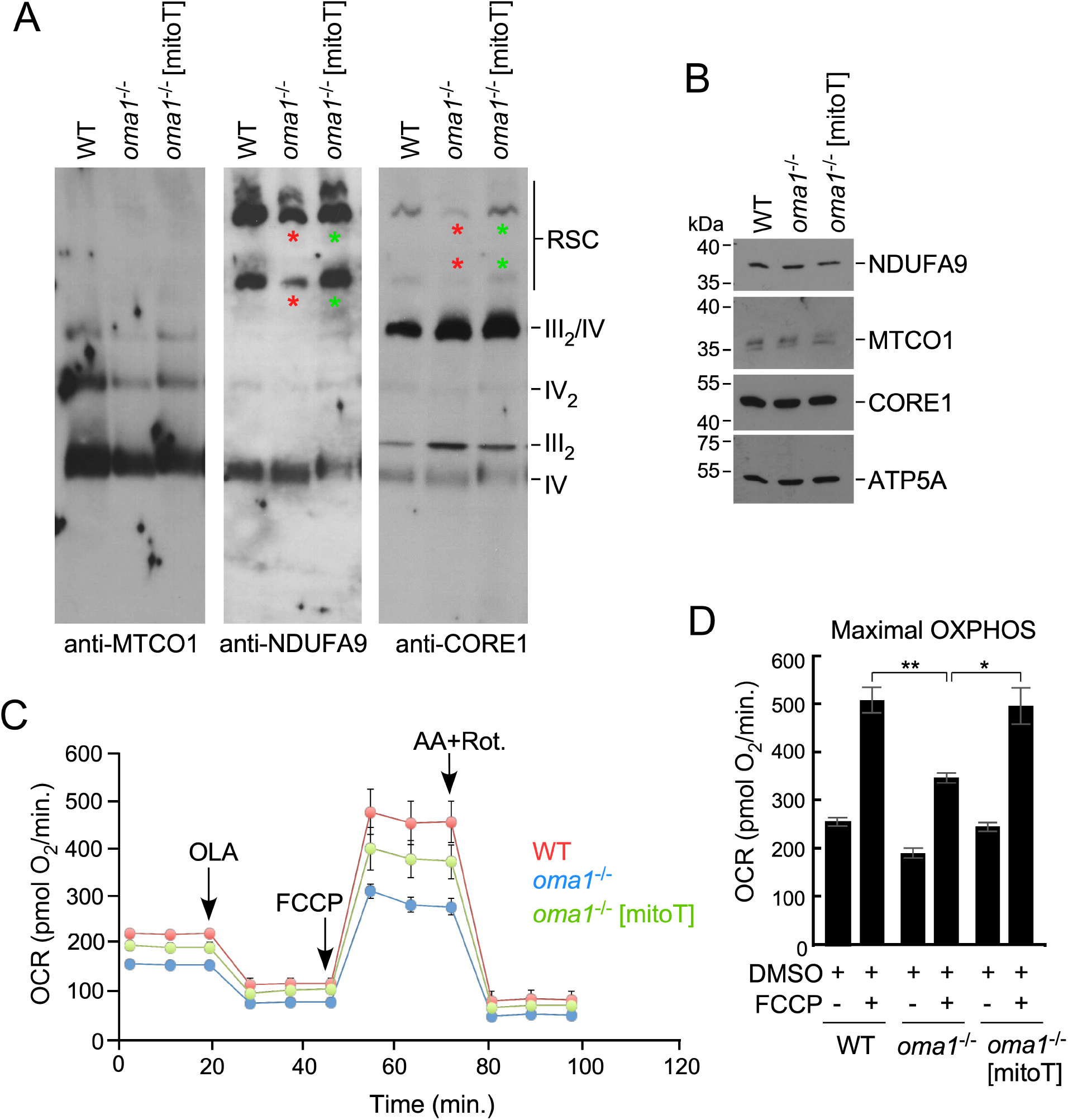
MitoT expression stabilizes respiratory complexes and mitigates bioenergetic deficit in OMA1-deficient cells. **(A)** Mitochondrial lysates from the respective cells were subjected to native gel electrophoresis and analyzed by western blot with indicated antibodies. Red asterisks highlight more labile respiratory supercomplexes, and green asterisks highlight stabilized respiratory supercomplexes. **(B)** Steady-state levels immunoblot analysis with indicated antibodies of mitochondrial lysates from the same samples as in A. **(C and D)** *In vivo* oxygen consumption rates (OCR) of indicated cell lines (50,000 cells/well) under basal, oligomycin A (OLA), FCCP and antimycin A + rotenone stimulated conditions were assessed by extracellular flux analysis. Cells were cultured in medium containing 10 mM galactose. Representative OCR graph is shown in C. Panel D focuses on FCCP-stimulated maximal respiration. Error bars, mean ± S.E. of n=3 biological replicates; \**p*<0.05, \*\**p*<0.01 by *t*-test.

We next evaluated the physiological relevance of mitoT-mediated RSCs stabilization by measuring *in vivo* oxygen consumption rates (OCR) in WT, *oma1*^−/−^ and *oma1*^−/−^ [mitoT] cells cultured in galactose. This culture condition forces cells to rely on oxidative metabolism (Gohil et al., 2010), and OMA1-deficient MEFs exhibit signs of bioenergetic deficit when cultured in galactose-supplemented medium (Bohovych et al., 2015). This condition does not significantly alter viability of these cells (Supplementary Fig. S5A). In line with our previous report (Bohovych et al., 2015), we found that oma*1*^−/−^ MEFs were unable to maximize their bioenergetic output in response to CCCP-treatment (Fig. 5C, D). In contrast, the mitoT-expressing oma*1*^−/−^ cells exhibited a nearly-WT bioenergetic profile - most notably being able to efficiently maximize their OCR in response to mild uncoupling (Fig. 5C, D). The observed effect does not appear to be merely due to an excess of the L-OPA1 form, as expression of the L-OPA1ΔS1 variant lacking OMA1 cleavage site (Mishra et al., 2014; Lee et al., 2017) in *oma1*^−/−^ MEFs did not mitigate bioenergetic deficit in these cells (Supplementary Fig. S5B and C). In fact, we found that such genetic manipulation reduced basal OCR of *oma1*^−/−^ cells (Supplementary Fig. S5C). These data indicate an additional mechanism is at play in OMA1-mediated regulation of bioenergetic output.

Finally, we used tetramethylrhodamine (TMRM) staining to examine basal mitochondrial membrane potential in the respective cells. We found that OMA1-depleted cells exhibited lower TMRM fluorescence thereby reflecting decreased mitochondrial membrane potential (Fig. 6A). Interestingly, mitoT expression did not improve mitochondrial polarization in these cells, indicating that membrane tethering via mitoT may not fully replace some functions related to mitochondrial ultrastructure and/or MICOS complex that are compromised in OMA1-deficient MEFs.

**Figure 6.**
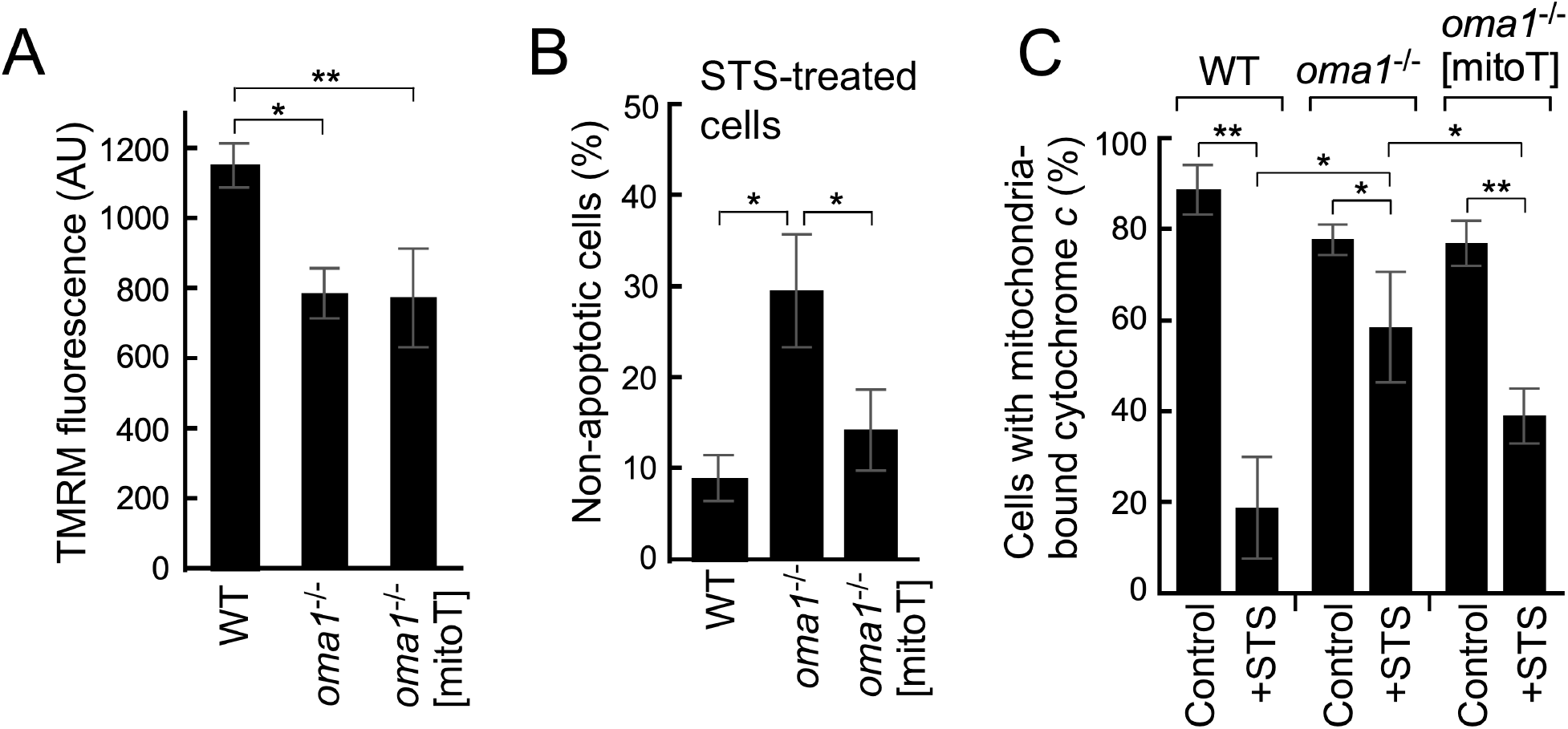
MitoT expression ablates apoptotic resistance in *oma1*^*−/−*^ cells. **(A)** Mitochondrial membrane potential in the indicated cell lines was assessed by flow cytometry analysis of tetramethylrhodamine (TMRM)-stained cells. Error bars, mean ± S.D. of n=3 biological replicates; \**p*<0.05, \*\**p*<0.01, by *t*-test. **(B, C)** Flow cytometry analysis of indicated cell lines that were treated with or without 1 mM staurosporin to assess the percentage of viable/apoptosis-resistant (Q3: 7-AAD negative, Annexin V-APC negative) cells (B) or cells with mitochondria-retained cytochrome *c* (C). Error bars, mean ± S.E. of n=3 biological replicates; \**p*<0.05, \*\**p*<0.01, by *t*-test.

Collectively, these results suggest that impaired RSCs stability and bioenergetic deficit in OMA1-deficient cells arise from impaired OM-IM contacts and altered mitochondrial ultrastructure.

### Anti-apoptotic effect of OMA1 deletion is linked to impaired membrane contacts

Previous reports established that loss of OMA1 renders cells resistant to apoptotic stimuli and attributed this effect to attenuated mitochondrial fission and cytochrome *c* release due to impaired L-OPA1 cleavage (Jiang et al., 2014; Varanita et al., 2015; Korwitz et al., 2016). Our findings prompted us to revisit this model and test whether impaired OM-IM connectivity may also be contributing to an anti-apoptotic effect in *oma1*^−/−^ cells. We thus examined Annexin V presentation and cytochrome *c* release – two key steps in the commencement of a classical apoptotic program – in WT, *oma1*^−/−^ and *oma1*^−/−^ [mitoT] MEFs treated with the apoptosis-inducing drug staurosporin. In line with previous reports (Jiang et al., 2014; Varanita et al., 2015; Korwitz et al., 2016), we observed that *oma1*^−/−^ cells accumulated significantly less externalized Annexin V, relative to WT control (Fig. 6B and Supplementary Fig. S6A). Likewise, staurosporin-treated *oma1*^−/−^ MEFs displayed significant impairment in cytochrome *c* release as compared to WT cells (Supplementary Fig. S3B and Fig. 6C). Remarkably, mitoT expression partially restored both staurosporin-induced Annexin V presentation and cytochrome *c* release in *oma1*^−/−^ MEFs (Fig. 6B and C). The expression of mitoT had no appreciable effect on mitochondrial fragmentation in the cells of interest (Supplementary Fig. S7). We conclude that reversal of apoptotic resistance in *oma1*^−/−^ [mitoT] cells is due to stabilized IM-OM contacts rather than enhanced mitochondrial fragmentation.

These findings further suggest that OM-IM connectivity is important for stress-associated membrane remodeling and initiation of the apoptotic program. Moreover, our data expand the current model of how OMA1 inactivation may relate to cellular apoptotic resistance and warrant further investigation of this issue.

## DISCUSSION

Remodeling of mitochondrial membranes is now being recognized as an integral part of multiple physiological processes in eukaryotic cells. Although the key players involved in sculpturing and dynamic rearrangement of the OM and IM have been identified, their coordinated function and the factors that orchestrate these mechanisms remain largely unclear. Here, we show that IM-anchored metallopeptidase OMA1 is dynamically associated with the MICOS complex, thereby stabilizing IM-OM contact sites and facilitating their remodeling in response to homeostatic insults. Mechanistically, this association appears to be mediated by OMA1’s interaction with the MIC60 subunit of MICOS, which we identified by proximity labeling, direct co-purification and co-fractionation analyses. Interestingly, similar to the recently reported MIC60-OPA1 association (Glytsou et al., 2016; Barrera et al., 2016), this interaction has previously eluded identification by the recent high-throughput proteomic analysis of MICOS (Guarani et al., 2015; Schweppe et al., 2017), probably due to the low cellular abundance of OMA1. Of note, we have also observed limited interaction between MIC60 and OPA1. However, the results of co-purifications in *opa1*^−/−^ and *oma1*^−/−^ MEF cells indicate that OMA1 and OPA1 associate with MICOS independently of one another.

What is the effect of OMA1 loss with respect to MICOS? Consistent with previous reports (Quiros et al., 2012; Anand et al., 2014; Korwitz et al., 2016), we observed that OMA1 deletion in MEFs did not result in any appreciable mitochondrial morphology under basal conditions. As expected, unlike the WT cells, *oma1*^−/−^ MEFs did not exhibit fragmented mitochondrial networks upon mild uncoupling due to impaired L-OPA1 processing and preservation of OPA1 oligomers. We found, however, that stressed OMA1-deficient mitochondria tended to form tubules that were morphologically distinct from those observed in unperturbed cells. Further analyses revealed that high-mass MICOS complexes were less stable under native gel electrophoresis conditions in *oma1*^−/−^ mitochondria even if cells were not challenged with an uncoupler. Accumulation of lower-mass, MIC27-containing complexes (Supplementary Fig. S2) likely reflects impaired association of MIC60-MIC19 subcomplex with other MICOS subunits known as the MIC10 subcomplex. Our data indicate a general nature of this defect, but suggest it becomes physiologically significant upon conditions of mitochondrial stress. In line with this notion are the outcomes of our mitochondrial ultrastructure analysis. We reproducibly observed that about half of *oma1*^−/−^ MEF cells exhibited aberrant mitochondrial ultrastructure; this might be in part due to harsh EM sample preparation conditions, which we believe have helped to reveal this subtle phenotype. More importantly, mitochondria in uncoupled *oma1*^−/−^ MEFs displayed gross ultrastructure alterations wherein the majority of cristae appeared to be dislocated from the OM and formed vesicle- or onion ring-like structures. Remarkably, the latter alterations strongly resemble ultrastructure defects observed in MICOS-depleted cells (Plecita-Hlavata et al., 2016; Glytsou et al., 2016 and Supplementary Fig. S3). Altogether, these data suggest that OMA1-MICOS association is required for efficient ICM and/or IBM rearrangement in response to homeostatic insults. Although neither MIC60 nor other core MICOS subunits seem to be a *bona fide* substrate of OMA1, and the exact mechanistic aspect of this association awaits an in-depth investigation, our results indicate that in the absence of OMA1, MICOS complex appears to be more labile and thus impeded in its ability to efficiently reorganize OM-IM contact sites in response to homeostatic insults. Such a model is consistent with our finding that expression of the OM-IM bridging mitoT tether substantially ameliorated uncoupling-induced ultrastructure abnormalities in *oma1*^−/−^ cells. A possibility exists that some of the above effects might also be connected to the inability of *oma1*^−/−^ mitochondria to rapidly produce S-OPA1. However, previous reports showing that L-OPA1 alone is sufficient to mediate mitochondrial dynamics and ultrastructure (Ishihara et al., 2006; Tondera et al., 2009; Anand et al., 2014; Lee et al., 2017) and that cells expressing a non-cleavable L-OPA1 variant do not appear to exhibit vesicular cristae (Lee et al., 2017) argue against this scenario.

What is the physiological role of OMA1-MICOS association? Results of our experiments with the mitoT membrane tether indicate this association is required for stabilization of respiratory supercomplexes and optimal respiratory output in response to physiological challenges. These data are consistent with a model wherein remodeling of both IBM and ICM ultrastructure and the maintenance of OM-IM connectivity are required to reorganize and/or stabilize respiratory complexes, thereby optimizing their performance to meet changing bioenergetic demands in response to physiological or stress stimuli (Mishra et al., 2014; Patten et al., 2014; Lee et al., 2017). Alternatively, OM-IM contacts may facilitate functions of certain carrier proteins, ultimately providing better substrate accessibility for RSCs. Our results also offer a plausible explanation as to how loss of OMA1 could impair RSCs stability and cause bioenergetic deficit in response to mild uncoupling (Bohovych et al., 2015; Quiros et al., 2012). These models are in line with the previously reported bioenergetic deficit in MICOS mutant cells (Darshi et al., 2012; Ott et al., 2012; Ott et al., 2015; Ding et al., 2015; Genin et al., 2016; Guarani et al., 2015).

Finally, our findings indicate that OMA1-MICOS mediated intermembrane connectivity is important for commencement of the apoptotic program. Earlier reports established that preservation of L-OPA1 oligomers via L-OPA1 overexpression (Varanita et al., 2015) or OMA1 depletion (Quiros et al., 2012; Jiang et al., 2014; Korwitz et al., 2016; Wai et al., 2016) results in an anti-apoptotic effect, presumably due to the maintenance of tight cristae junctions and subsequent obstruction of cytochrome *c* release from the intercristae space in response to apoptotic stimuli (Frezza et al., 2006; Yamaguchi et al., 2008; Varanita et al., 2015). Our data expand this model of apoptotic resistance of *oma1*^−/−^ cells as follows. First, we observed that physiological insults to OMA1-deficient cells promoted formation of OM-disconnected yet locked cristae that are likely to encapsulate and retain the majority of available cytochrome *c* pools. Indeed, these results were consistent with the elevated resistance of *oma1*^−/−^ MEFs to the apoptosis-inducing drug staurosporin, as indicated by the significant decrease in Annexin V-stained cells and enhanced retention of mitochondria-bound cytochrome *c*. Second, we found that enforcing OM-IM contacts in OMA1-deficient MEFs through mitoT tether expression was sufficient to curtail the cells’ resistance to staurosporin-induced apoptosis. In line with this finding we observed that mitoT-expressing *oma1*^−/−^ MEFs exhibited increased cytochrome *c* release upon staurosporin treatment as compared to untransformed cells. Collectively, these results suggested an updated model wherein not only cristae junction tightness, but also IM-OM connectivity is important for molecular events preceding apoptosis initiation. Overall, these data further our mechanistic understanding of OMA1’s role in mitochondrial architecture, bioenergetics and cell death.

While this paper was in preparation, a report appeared demonstrating that MIC60 and MIC19 MICOS subunits are connected to the SAM50 core component of the OM sorting and assembly machinery (SAM) complex via the MIC19 subunit to dynamically shape cristae architecture (Tang et al., 2020). Tang et al. showed that under certain homeostasis-challenging conditions, OMA1 protease can mediate the N-terminal cleavage of MIC19, thereby destabilizing SAM-MICOS association and impacting mitochondrial architecture. Although these data are generally in line with our results described in the present study, we were unable to obtain any evidence for OMA1-mediated MIC19 cleavage, likely due to differences in sample isolation procedures (whole cell lysates vs. purified mitochondria) and/or types of antibodies and cell lines used. Therefore, the exciting new mechanism by which MICOS is dynamically regulated by OMA1 awaits further mechanistic investigation.

## ACKNOWLEDGEMENTS

We thank Nataliya Zahayko, Colton Roesner and Drew Harrahill for technical assistance, and Jonathan Dietz and Dr. Iryna Bohovych for their help with gradient fractionation and native gel electrophoresis analyses; we thank Dr. Jennifer L. Fox for editorial help. Mass spectrometry analysis was performed at the Metabolomics & Proteomics Core Facility at the University of Nebraska-Lincoln, which is supported by NIH grant P30 GM103335. We also acknowledge help form the Flow Cytometry Service Center and the Morrison Microscopy Core Facility at the University of Nebraska-Lincoln. This work was supported by NIH grants GM108975, GM131701-01 (O.K.) and P30 GM103335 (O.K. through the Nebraska Redox Biology Center), and the Deutsche Forschungsgemeinschaft (DFG) CRC 1218 – project-ID 267205415 – project B12 (A.S.R.).

## AUTHOR CONTRIBUTIONS

R.M.L., M.P.V., and O.K. designed experiments. R.M.L., and O.K. wrote the paper with the input from all authors. R.M.L., and M.P.V. conducted the experiments and analyzed data. R.A. and A.S.R. provided critical data and reagents and helped with data interpretation.

## STAR METHODS

Detailed methods are provided in the online version of this paper and include the following.

### RESOURCE AVAILABILITY

#### Lead Contact

All requests for additional information, resources and reagents should be directed to the Lead Contact, Dr. Oleh Khalimonchuk.

#### Materials Availability

All reagents generated in this study are available from the Lead Contact upon request and with completed materials transfer agreement.

### EXPERIMENTAL MODEL AND SUBJECT DETAILS

#### Cell Lines and Cell Culture Conditions

Wild type mouse embryonic fibroblasts and isogenic MEF *oma1*^−/−^ cell lines were a kind gift from C. Lopez-Otin (Univ. of Oviedo). We have also obtained another set of WT and *opa1*^−/−^ MEF cells from ATCC (CRL-2991 and CRL-2995 respectively). Human embryonic kidney HEK293T cells (ATCC CRL-3216) were a gift from Dr. Sathish Natarajan (Univ. of Nebraska-Lincoln). MEF cells were cultured in DMEM medium with addition of 10% fetal calf serum (Atlanta Biologicals), 10 mM glucose, 2 mM pyruvate, 4 mM L-glutamine, 50 U/ml penicillin, with addition of 0.1% β-mercaptoethanol (all from Thermo Fisher Scientific). HEK293T cells were cultured in the same medium, albeit without the addition of β-mercaptoethanol. All cells were cultured in a humidified CO_2_ incubator at 37°C, 95% air and 5% CO_2_ mixture. Cells were trypsinized for 5 min in 0.05% trypsin after a wash with Ca^2+^- and Mg^2+^-free PBS. Cells were counted using Countess Cell Counter and Countess Cell Counting Slides (Invitrogen). Cell lines were regularly analyzed for mycoplasma contamination.

Transient transfections were performed using Lipofectamine 3000 reagent (Thermo Fisher Scientific) according to the manufacturer-supplied protocol using the high range of the recommended Lipofectamine 3000 amounts (3 μl per well for 12-well plates, 7.5 μl per well for 6-well plates, and 44 μl per 10 cm^2^ dish). Experiments with transfected cell lines were conducted 24 h post-transfection.

#### Key Reagents

Tables listing antibodies, media, chemicals and kits used in this study can be found in Key Resources Table.

### METHOD DETAILS

#### Generation of MitoT-expressing Cell Lines

The GFP-tagged mitoT construct was generated by fusing the N-terminal portion of SCO1 (amino acid residues 1-116), a 12 residues-long unstructured linker sequence derived from *E. coli* LacI, and the N-terminal region of TOM20 (amino acid residues 1-20) followed by a 6xHis tag. This chimera was generated by an overlap extension PCR using the following oligonucleotides:

5’-AAAGGATCCATGGCGATGCTGGTCCTAGTACCC-3’ (F-SCO1-BamHI)
5’-GACATCGTATAACGTTACTGGTTTCTTGACGTGCTTCATTCCAGC-3’ (R-SCO1-LacI)
5’-AGTAACGTTATACGATGTCGCAGAGATGGTGGGTCGGAACAGCGC-3’ (F-LacI-TOM20)
5’-CAAGAATTCGTGATGGTGATGGTGATGGAAGTAGATGCAGTACCCA-3’ (R-TOM20-His-EcoRI)

The resulting 468-bp product harboring BamHI and EcoRI restriction sites was cloned into the pcDNA3-eGFP vector (Addgene #13031). The resulting plasmid was validated by DNA sequencing, purified, and transfected into WT or *oma1*^−/−^ MEF cells, and following 24 hours after transfection selected in 3 μg/ml puromycin for 1 week. The transfected cell population was subjected to discriminatory sorting based on GFP signal versus mock-transfected cells on FACS Aria II (BD Biosciences), and the resulting GFP-positive cells were further selected for 1 week in a culture with 3 μg/ml puromycin, resulting in a cell line stably expressing mitoT-GFP. Confocal microscopy and western blot analyses confirmed that the expression of mitoT-GFP did not change after 1 month in culture.

#### Flow Cytometry Analyses

Flow cytometry was performed either on FACSCalibur II (BD Biosciences) or Cytek DxP10 (Cytek Biosciences) flow cytometers recording a minimum of 10,000 events per sample, with analysis of the resulting data using FlowJo 10 software (FlowJo Inc).

Mitochondrial membrane potentials were determined using TMRM dye staining according to a previously described method (Rodriguez-Rocha et al., 2013). In brief, cells in confluent wells in a 12-well cell culture plate were subjected to either 2 μM CCCP or mock for 1 hour, trypsinized, washed twice with PBS, resuspended in 1 ml of PBS, and stained with 50 nM TMRM for 15 minutes on ice. Cells were then pelleted at 400 x*g*, resuspended in fresh PBS, and measured on a Cytek DxP10 flow cytometer within 30 minutes. TMRM fluorescence was measured using 561 nm excitation, and 580/20 nm emission filters.

Annexin V presentation and cell viability were assessed after an 8-hour exposure to 1 μM staurosporin or mock-treatment using 7-AAD APC-Annexin Kit (BioLegend) according to manufacturer’s recommendations.

Cytochrome *c* release was assessed after an 8-hour exposure to 1 mM staurosporin or mock using a cytochrome *c* release kit (EMD Millipore) per manufacturer’s recommendations. Upon necessity to avoid spectral overlap with the GFP fluorescence, the FITC-labeled anti-cytochrome *c* antibody supplied in the kit was substituted with Alexa Fluor 647-labeled anti-cytochrome *c* antibody (BioLegend).

#### Imaging Techniques

For confocal microscopy imaging, cells were stained in 1 cm glass bottom dishes (MatTek Corp.) with 5 nM Hoechst 33342 (Thermo Fisher Scientific) and/or 5 μM Mitotracker DeepRed (Thermo Fisher Scientific) dyes in Phenol Red-free DMEM medium for 20 minutes in the dark, washed 2x with PBS, and supplemented with 2 ml of Phenol Red-free DMEM medium. The imaging was conducted using a Nikon A1R-Ti2 confocal system (Nikon, Japan). Multicolor images were generated using ImageJ software (NIH) by merging appropriate channels after assigning colors to each of them (green for GFP channel, blue for Hoechst 33342, and red for MitoTracker DeepRed).

For TEM, cells exposed to 2 μM CCCP or mock for 1 h were trypsinized, washed 3x with PBS, fixed in 100 mM sodium cacodylate buffer pH=7.4 with 2.5% glutaraldehyde, and processed as previously described (Graham and Orenstein, 2007). TEM was performed on a Hitachi H7500 TEM microscope. Pictures were taken at 10,000x and 25,000x magnification from three different grids per sample. Mitochondria on each photo were assessed for ultrastructure type, which was classified as abnormal/vesicular or normal. The number of mitochondria of each type was counted in each grid, and percent of total for each type was calculated in each sample.

#### Biochemical Assays

For cell lysis preparation 2 million cells per sample were lysed by tumbling at 4°C for 2 hours in NP-40 lysis buffer containing 1x protease inhibitors cocktail and 2 mM PMSF freshly added prior to the lysis. The lysates were supplemented with 6x Laemmli buffer containing 5% β-mercaptoethanol and after boiling for 5 min at 100°C were loaded into a 10% polyacrylamide gel for subsequent immunoblot analysis. For IP, cells were grown to 90% confluence, and 24 hours after transfection were trypsinized, washed 3x with PBS, and lysed. Anti-Myc magnetic beads (Thermo Fisher Scientific, 88842), anti-GFP magnetic beads (MBL International, D153-9), and isotype IgG magnetic beads (MBL International, M076-11) were used with the lysates according to manufacturer’s recommendations. Sucrose gradient separation of mitochondrial proteins was performed as previously described (Khalimonchuk et al., 2010) and analyzed by western blotting with relevant antibodies. Mitochondria (200 μg) were lysed using 20 μM HEPES buffer pH 7.4 with 100 mM NaCl and 1% digitonin on ice for 15 minutes. The lysate was centrifuged at 20000 *x g* for 10 min. at 4°C. 150 μl of the supernatant was added to 40 μl of anti-Mic60 agarose beads containing 1500 μl of 20 μM HEPES buffer pH 7.4 with 10 0mM NaCl and 0.1% digitonin and was kept rotating at 4°C for 16h. Proteins were eluted from the beads using 2x Laemilli buffer without any reducing agent.

For blue native gel electrophoresis of mammalian mitochondria, Native PAGE Novex Bis-Tris Gel System and Native PAGE Novex 3-12% Bis-Tris Gels (Thermo Fisher Scientific) were used along with other reagents according to manufacturer’s recommendations for organelle protocols. Mitochondria were extracted from MEF cells as described previously (Chen et al., 2012), solubilized in a sample buffer containing 0.5% digitonin and used as samples according to the manufacturer’s protocol. Proteins from the resulting gels were transferred to PVDF membranes (Thermo Fisher Scientific) and analyzed by western blotting with relevant antibodies.

#### Immunoblotting

Proteins were detected using the primary antibodies indicated in the Key Resources Table. Relevant protein bands were visualized using secondary horseradish peroxidase-coupled secondary goat-anti-mouse (Jackson ImmunoResearch) and goat-anti-rabbit (Cell Signaling Technologies) secondary antibodies and chemiluminescent reagents (Thermo Fisher Scientific).

#### BioID Analysis

The pcDNA3.1-mycBioID plasmid was a kind gift from K. Roux (Addgene #35700). Human *OMA1* and human *SURF1* (matched negative control) were amplified using Phusion High Fidelity DNA Polymerase (Thermo Fisher Scientific) from HEK293T cDNA library and subcloned into the plasmid using NheI and EcoRI restriction sites to create BirA fusion proteins. The plasmids were expressed in HEK293T cells, and proximity interactions were analyzed according to a previously described protocol (Roux et al., 2013). Briefly, 48 hours after transfection, 20 million transfected cells were lysed and tumbled overnight with Streptavidin magnetic beads (Thermo Fisher Scientific), after which the beads were washed 3x with lysis buffer, boiled, and subjected to SDS-PAGE. Abundant protein bands were extracted out of the gel and subjected to mass spectrometry detection on a QSTAR XL Hybrid LC/MS/MS (Applied Biosystems). Results were analyzed using Scaffold 4 (Proteome Software).

#### Extracellular Flux Analysis

The cells were seeded in parallel in two identical Seahorse cell culture plates (Agilent Technologies), one of which was used for the actual experiment, while the other was used for cell count for normalization purposes. Extracellular Flux experiments were performed on a Seahorse XFe24 Analyzer (Agilent Technologies) using XF Cell Mito Stress Test and XF Glycolysis Stress Test kits (Agilent Technologies) per manufacturer’s protocols. Data were analyzed using manufacturer’s calculation templates for Microsoft Excel 365. Summary bar graphs were plotted in Microsoft Excel 365 based on the data from the original analysis templates.

### QUANTIFICATION AND STATISTICAL ANALYSIS

Unless stated otherwise, the data were analyzed in Microsoft Excel 365 using the Data Analysis package. All data are shown as mean ± S.D., unless indicated differently. *P* values were calculated using paired two-tailed *t*-tests when analyzing paired samples and unpaired two-tailed *t*-tests when analyzing unpaired samples. *P* values less than 0.05 were considered significant. Violin graphs were plotted in GraphPad Prism 4.

For image quantification, images on X-ray films were digitalized and analyzed using Image J software.

For BioID LC-MS/MS data analysis, the following stringency criteria for candidate protein acceptance were applied. First, common background proteins were discarded. Second, candidate proteins with less than three spectral counts were considered as low-confidence hits and not pursued further. Third, candidate hits were checked against the CRAPome contaminant repository database (https://www.crapome.org) and unrelated mitochondrial control (SURF1) BioID analysis data to further eliminate nonspecific binding partners.

## SUPPLEMENTAL INFORMATION

Supplemental information includes seven Supplemental Figures and can be found with this article online.

---

KEY RESOURCES TABLE.

KEY RESOURCES TABLE
| REAGENT / RESOURCE | SOURCE | IDENTIFIER |
| --- | --- | --- |
| *Antibodies* |  |  |
| OMA1 | Santa Cruz | Cat.# sc-515788 |
| OPA1 | BD Biosciences | Cat.# 612606 |
| OPA1 | Novus Biologicals | Cat.# NB110-5529 |
| MIC60 | Protein Tech | Cat.# 10179-1-AP |
| MIC27 | Aviva Systems Biology | Cat.# OAAF03292 |
| MIC19 | Protein Tech | Cat.# 10179-1-AP |
| MIC10 | Aviva Systems Biology | Cat.# ARP44801_P050 |
| MIC13 | Proteintech | Cat.# 25514-1-AP |
| VDAC | Abcam | Cat.# ab14734 |
| HSPD1 | Santa Cruz | Cat.# sc-13966 |
| YME1L | Protein Tech | Cat.# 11510-1-AP |
| GAPDH | Protein Tech | Cat.# 60004-1-Ig |
| GAPDH | GeneTex | Cat.# GTX627408 |
| NDUFA9 | Abcam | Cat.# ab14713 |
| CORE1 | Abcam | Cat.# ab110252 |
| ATP5A | Abcam | Cat.# ab14748 |
| AFG3L2 | Abcam | Cat.# ab68023 |
| AFG3L2 | Aviva Systems Biology | Cat.# ARP4678055 |
| MTCO1 | Abcam | Cat.# ab14705 |
| GFP | Invitrogen | Cat.# a6455 |
| GFP | Santa Cruz | Cat.# sc-9996 |
| Myc | Roche | Cat.# 11667149001 |
| MFN1 | Abcam | Cat.# ab57602 |
| MFN2 | Abcam | Cat.# ab124773 |
| DRP1 | Santa Cruz | Cat.# sc-271583 |
| MIC60-agarose beads | Santa Cruz | Cat.# sc-390707 AC |
| Sterptavidin-magnetic beads | Thermo Fisher Scientific | Cat.# 88816 |
| Myc-magnetic beads | Thermo Fisher Scientific | Cat.# 88842 |
| GFP-magnetic beads | MBL International | Cat.# D153-9 |
| IgG-magnetic beads | MBL International | Cat.# M076-11 |
| 2° HRP-anti mouse | Jackson ImmunoResearch | Cat.# 115-035-068 |
| 2° HRP-anti rabbit | Cell Signaling Technologies | Cat.# 7074S |
| *Cell Lines* |  |  |
| HEK293T | ATCC | Cat.# CRL-3216 |
| Wild-type MEF | ATCC | Cat.# CRL-2991 |
| *opa1*<sup>-/-</sup> MEF | ATCC | Cat.# CRL-2995 |
| Wild-type MEF | Quiros *et al.*, 2012 | NA |
| *oma1*<sup>-/-</sup> MEF | Quiros *et al.*, 2012 | NA |
| Wild-type MEF [mitoT] | This study | NA |
| *oma1*<sup>-/-</sup> MEF [mitoT] | This study | NA |
| *Media, Chemicals and Reagents* |  |  |
| DMEM medium | Hyclone | Cat.# SH30243.01 |
| L-glutamine | Gibco | Cat.# 25030081 |
| Glutamax | Gibco | Cat.# 35050061 |
| Sodium Pyruvate | Gibco | Cat.# 11360070 |
| FBS, natural source | Hyclone | Cat.# SH30396.03 |
| Gentamicin | Gibco | Cat.# 15750060 |
| Penn/Strep mix | Hyclone | Cat.# 15070063 |
| Trypsin | Hyclone | Cat.# SH30236.02 |
| Lipofectamine 3000 | Gibco | Cat.# L3000015 |
| Opti-MEM I medium | Gibco | Cat.# 31985070 |
| FluoroBrite DMEM medium | Gibco | Cat.# A1896702 |
| Halt protease inhibitor | Thermo Fisher Scientific | Cat.# 87785 |
| Mitotracker DR-FM dye | Invitrogen | Cat.# M22426 |
| Hoechst 33342 dye | Invitrogen | Cat.# H3570 |
| Propidium Iodide | Invitrogen | Cat.# P3566 |
| Seahorse XF DMEM medium | Agilent | Cat.# 103575-100 |
| Digitonin | Sigma | Cat.# D141 |
| NP-40 | Sigma | Cat.# I3021 |
| NativePage 4-16% Bis-Tris precast gels | Invitrogen | Cat.# BN1002BOX |
| *Commercial Kits/Assays* |  |  |
| Super Signal West Dura Substrate | Thermo Fisher Scientific | Cat.# 34075 |
| XF Cell Mitostress Kit | Agilent | Cat.# 103015-100 |
| XFe24 FluxPack | Agilent | Cat.# 100867-100 |
| *Software* |  |  |
| Image J | NIH | https://imagej.nih.gov/ij/ |
| KaleidaGraph v.4.02 | Synergy Software | NA |
| GraphPad Prism 4 | GraphPad Software | NA |
| Scaffold 4 | Proteome Software | NA |
| FlowJo 10 | FlowJo Inc. | NA |
| Microsoft Excel 365 | Microsoft | NA |

