## Supplemental Information for "Protease OMA1 Modulates Mitochondrial Bioenergetics and Ultrastructure through Dynamic Association with MICOS Complex"

### Supplemental Figure Legends

**Figure S1, related to Figure 1. BioID analysis of OMA1 physical interactome. (A)** Top, Schematic depiction of OMA1-BirA construct used in BioID proximity labeling analysis. Bottom, List of relevant proteins identified by BioID proteomic analysis. **(B)** HEK293T cells were transfected with OMA1-GFP construct or mock transfected. Mitochondrial lysates from the respective cells were subjected to immunoprecipitation with anti-GFP magnetic beads. Precipitated proteins from the initial sample (load), unbound fraction (unbound), last wash (wash), and eluate were analyzed by western blot with indicated antibodies.

**Figure S2, related to Figure 2. Stability of high-mass MICOS complexes is compromised in *oma1*$^{-/-}$ cells. (A)** Schematic depiction of MICOS machinery and its partner protein complexes forming an IM-OM contact site. Some proteins are not shown for simplicity. **(B)** Mitochondrial lysates from WT and *oma1*$^{-/-}$ fibroblasts were analyzed by blue-native gel electrophoresis and immunoblotting with antibodies against indicated MICOS subunits as well as respiratory complex V subunit ATP5A (loading control).

**Figure S3, related to Figure 3. MICOS-deficient cells exhibit altered cristae morphology characterized by concentric membrane stacks and lack of cristae junctions.** Representative

TEM images of mitochondria in WT (left) and MIC13 knockout (right) HEK293T cells. Scale bars, 0.5 μm.

**Figure S4, related to Figure 4. Quantitative assessment of cell survival by flow cytometry. (A)** Viability over indicated periods of time of WT and *oma1*$^{-/-}$ fibroblasts with or without stable mitoT expression. Cells (500,000) were seeded per well in a 6-well plate. Growth was monitored for 3 days until over-confluence was reached. Events (10,000) were counted using Hoechst 33342 and propidium iodide staining parameters; healthy cells were PI-negative and Hoechst 33342-positive. Hoechst 33342-negative cells were not accounted for. Error bars, mean ± S.D. of n=3 biological replicates. **(B)** Steady-state levels of mitoT-GFP, OMA1, OPA1 and HSPD1 (loading control) in indicated cells that were incubated with or without 2 μM CCCP. Note that mitoT undergoes additional processing in the WT MEFs.

**Figure S5, related to Figure 5. Bioenergetic deficit in *oma1*$^{-/-}$ cells is not directly related to** **(A)** Steady-state levels immunoblot analysis of OMA1, YME1L, GAPDH and Myc-tagged OPA1 variant lacking S1 processing site (OPA1ΔS1) with relevant antibodies in mitochondrial lysates from indicated cells, with and without CCCP treatment. **(B)** Extracellular flux analysis of *in vivo* oxygen consumption rates in indicated MEF cells. Cells were cultured in medium containing 10 mM galactose, then transferred into standard Seahorse XF medium and analyzed at 50,000 cells/well under basal, oligomycin A (OLA), FCCP and antimycin A + rotenone stimulated conditions. Error bars, mean ± S.E. of n=4 biological replicates.

**Figure S6, related to Figure 6. Apoptotic resistance of *oma1*$^{-/-}$ cells. (A)** Quantitative assessment of cell survival by flow cytometry. Indicated cells were cultured in 10 mM glucose or 10 mM galactose-supplemented medium and stained with propidium iodide (PI). Healthy cells were identified as PI-negative. Data represents means ±S.E. of 3 biological replicates. **(B)** Flow cytometry analysis of healthy (Q3: 7-AAD negative, Annexin V-APC negative), early apoptotic (Q1: 7-AAD negative,

Annexin V-APC positive), late apoptotic (Q2: 7-AAD positive, Annexin V-APC positive) and necrotic (Q4: 7-AAD negative, Annexin V-APC positive) cells. Data are presented as mean ±S.E., n=3; *$p$<0.05, ***$p$<0.001 by *t*-test. **(C)** Left: Representative flow cytometry histograms assessing cytochrome *c* release in healthy (Control) and 1 mM staurosporin-treated (+STS) WT and *oma1*$^{-/-}$ fibroblasts. Iso, isotype control used to monitor antibody specificity. 50,000 cells were analyzed in each experiment. Right: Quantitative analysis of mitochondria-bound cytochrome *c* in the cells of interest. Data are presented as mean ±S.E. of 3 independent experiments; *$p$<0.05, **$p$<0.01, ***$p$<0.001 by *t*-test.

**Figure S7, related to Figure 6. Expression of mitoT does not induce mitochondrial fragmentation in *oma1*$^{-/-}$ cells.** Representative merged *in vivo* confocal microscopy images of mitochondrial network (visualized with mCherry, red signal) and mitoT-GFP (green signal) in indicated cells. Yellow color indicates signal overlap. Scale bar, 20 μm.

A

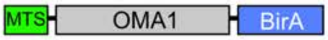


| Identified ORFs | Unique Peptide Sequences |
|---|---|
| ATP5B | 37 |
| ATP5A1 | 16 |
| ATAD3 | 17 |
| PHB1 | 3 |
| PHB2 | 4 |
| MIC60 | 24 |
| SLC25A6 | 17 |
| PRDX3 | 3 |

B

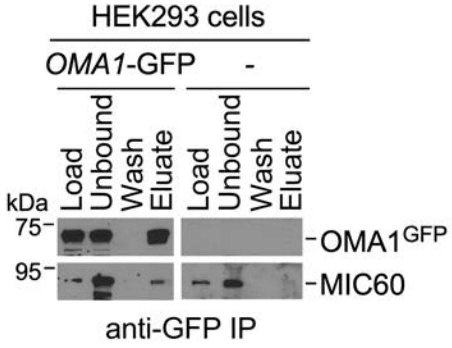


Supplementary figure S1

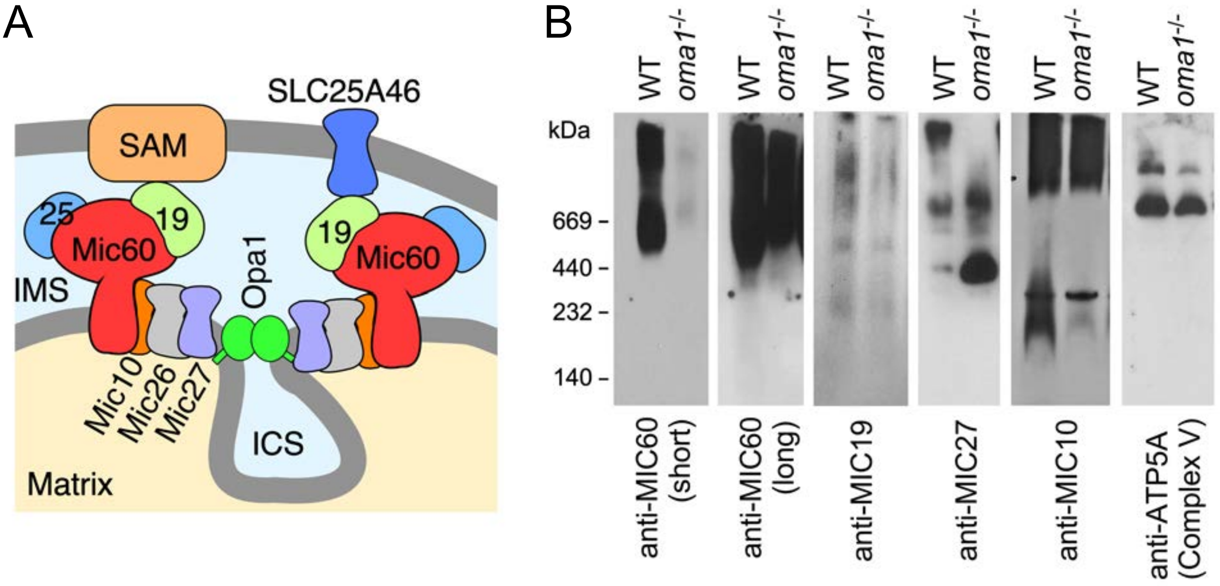


Supplementary figure S2

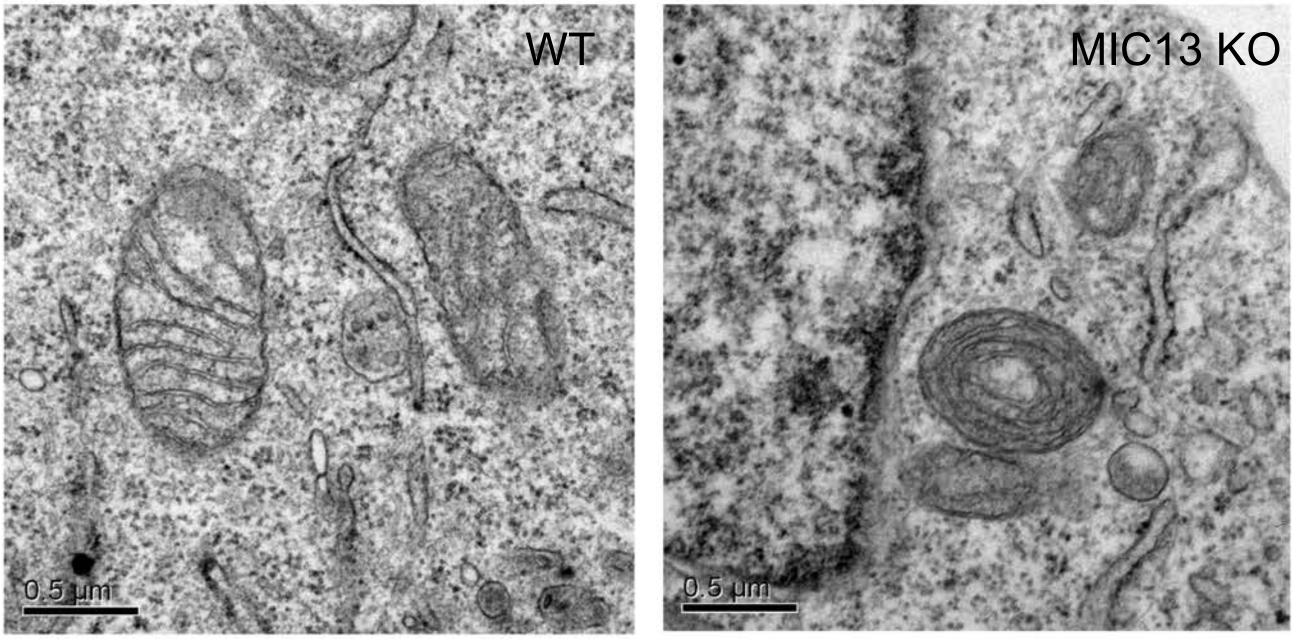


Supplementary figure S3

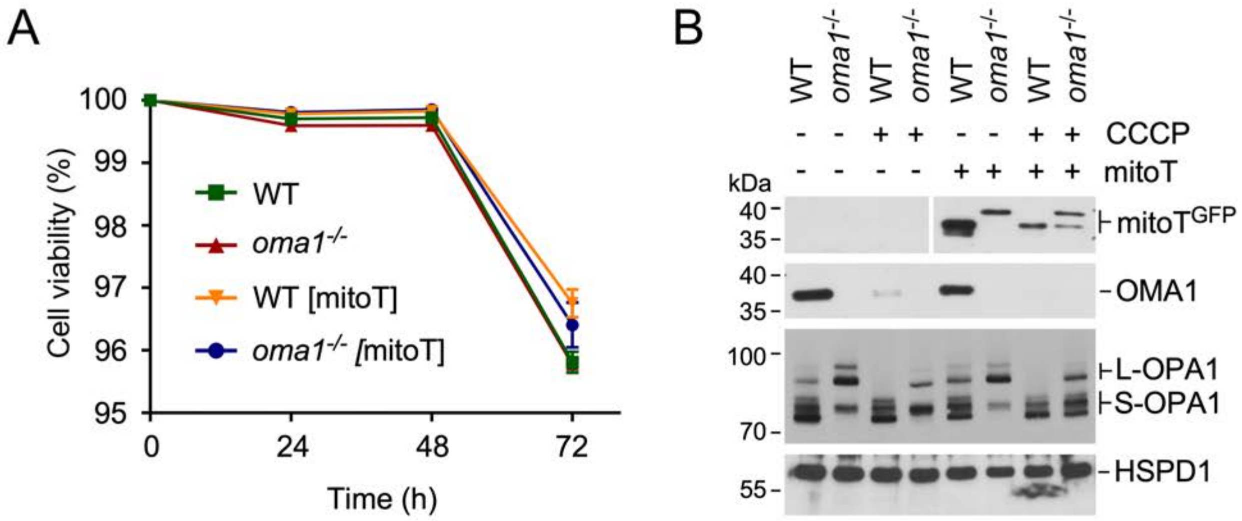


Supplementary figure S4

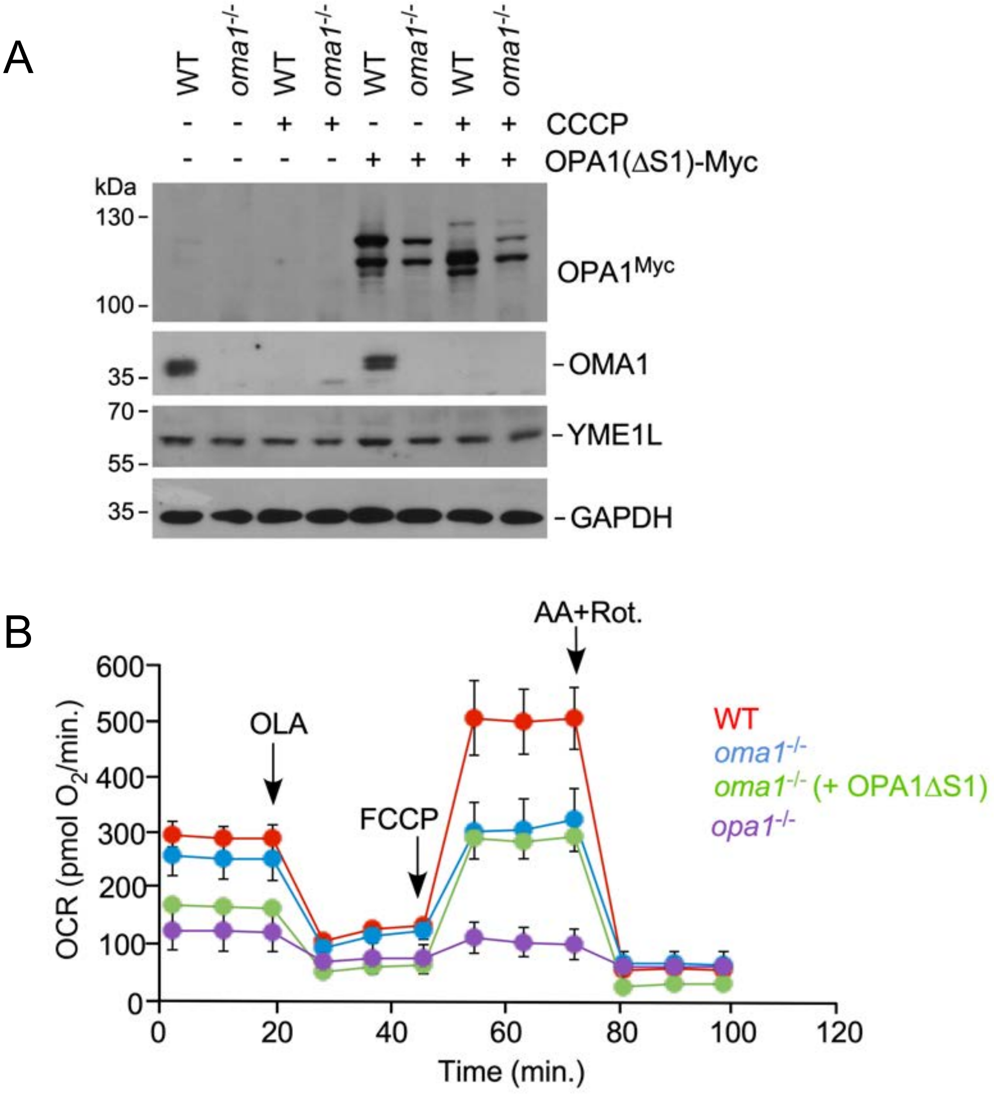


Supplementary figure S5

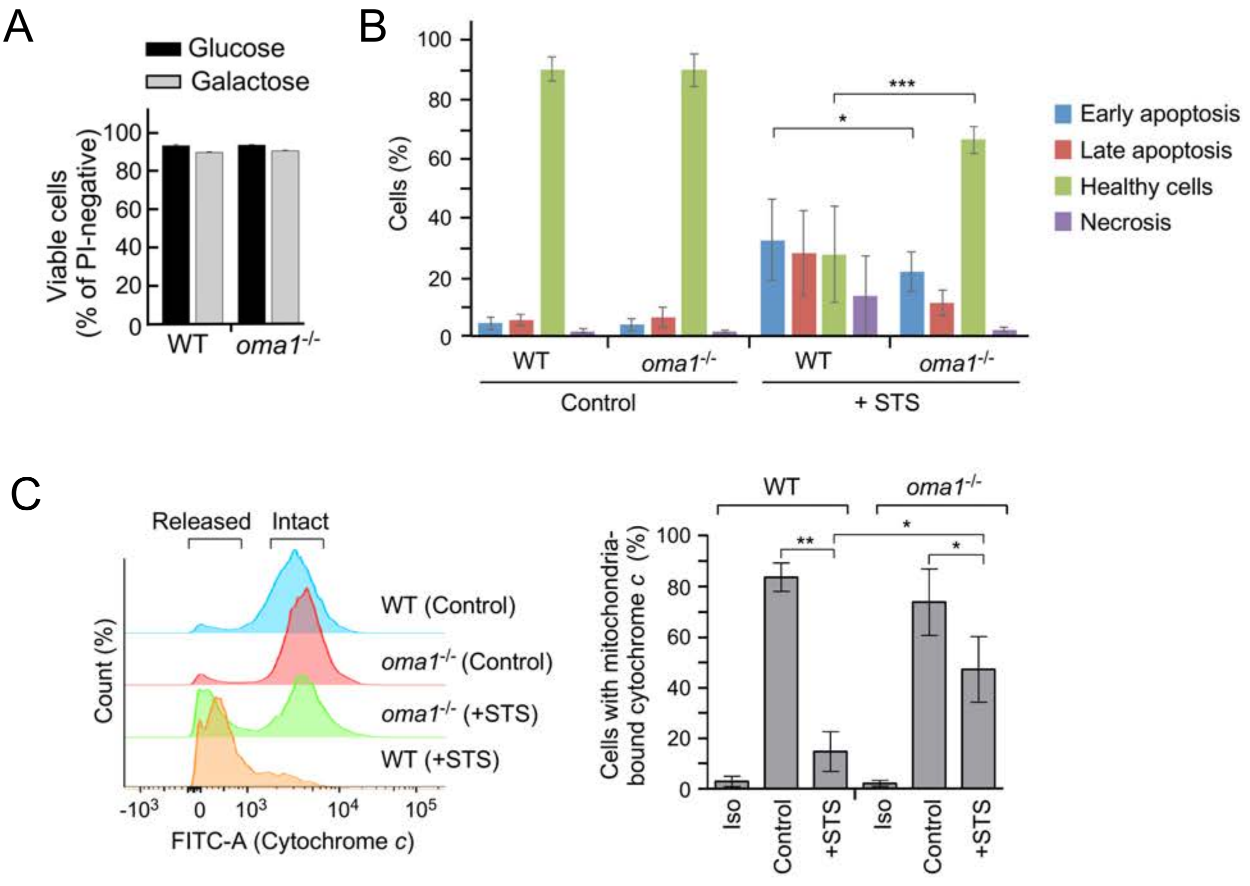


Supplementary figure S6

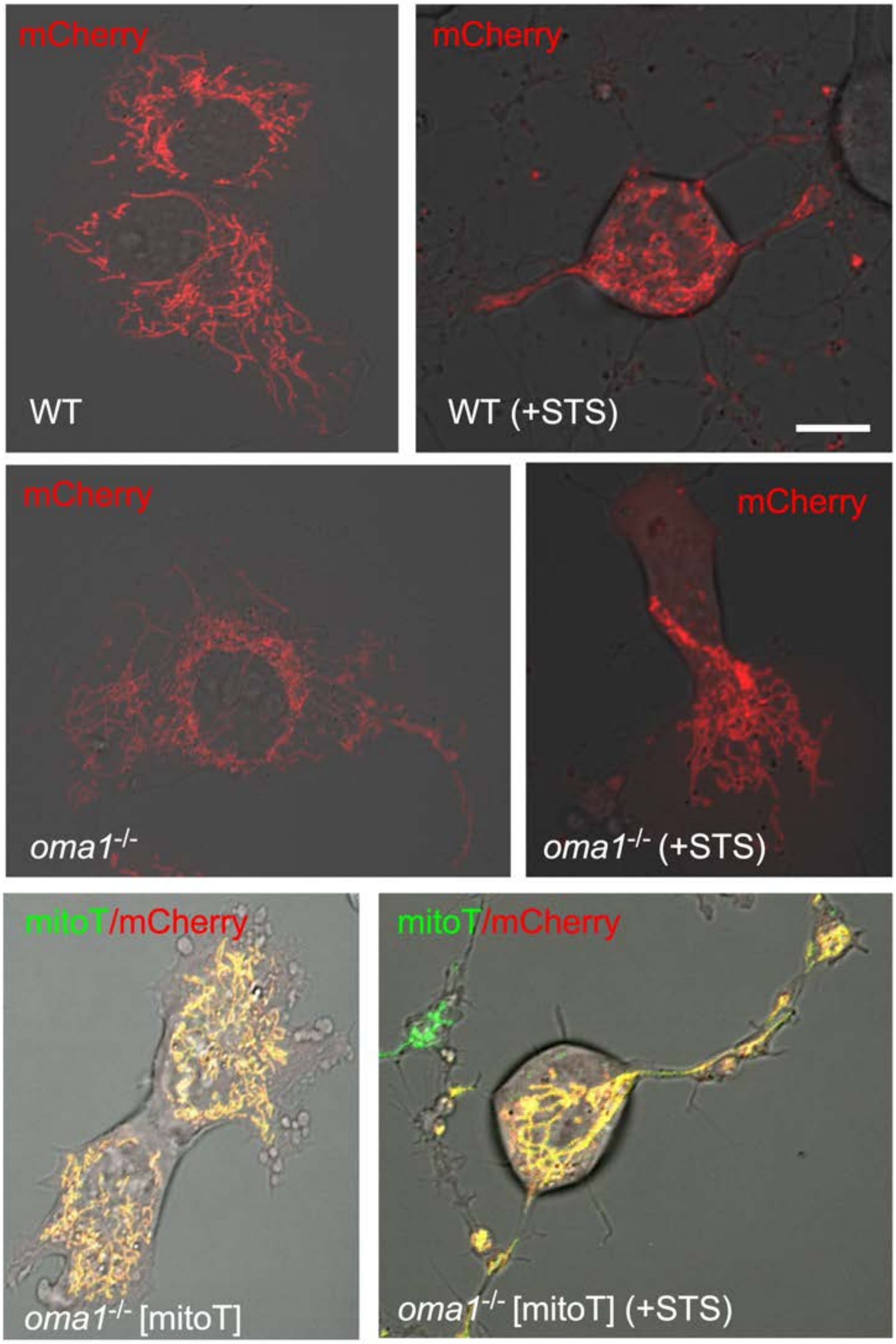


Supplementary figure S7
